# *Coxiella burnetii* strains elicit distinct inflammatory responses in human macrophages

**DOI:** 10.1101/2025.08.10.669536

**Authors:** Madhur Sachan, Amanda Dragan, Het Adhvaryu, Daniel E Voth, Rahul Raghavan

**Author notes:** Author for correspondence: Rahul Raghavan |.

## Abstract

*Coxiella burnetii*, the causative agent of human Q fever, subverts macrophage antimicrobial functions to establish an intracellular replicative niche. To better understand the host-pathogen interactions, we investigated the transcriptional responses of human alveolar macrophages (hAMs) infected with virulent (NMI, G), attenuated (NMII), and avirulent (Dugway) strains of *C. burnetii*. RNA sequencing analysis revealed that all strains activated proinflammatory pathways, particularly IL-17 signaling, though the magnitude and nature of the response varied by strain. Dugway infection induced the most robust transcriptional response and consistent M1-like macrophage polarization, while responses to NMI and NMII were more variable. Cytokine assays confirmed significant secretion of effectors downstream of IL-17 signaling, but only at later stages of infection. Single-cell RNA sequencing further revealed heterogeneity in macrophage response to *C. burnetii* infection, with distinct subpopulations exhibiting divergent inflammatory profiles. These findings highlight the complexity of macrophage responses to *C. burnetii* and underscore the importance of strain-specific and cell-specific factors in shaping host immunity. Understanding these dynamics may inform the development of targeted therapies for Q fever.

## INTRODUCTION

Alveolar macrophages typically provide the first line of defense against invading pathogens (1, 2). These monocyte-derived cells secrete cytokines to eliminate pathogens and to recruit other immune cells as part of the initial acute inflammatory response. One of the proinflammatory cytokines produced by immune cells in lungs is IL-17, which has a protective role against pulmonary intracellular pathogens such as *Mycobacterium tuberculosis* and *Legionella pneumophila* (3, 4). IL-17 binds to receptors on macrophage surface and triggers the production of antimicrobial peptides, secretion of chemokines and alteration of macrophage polarization towards the proinflammatory M1 phenotype, thereby accelerating pathogen destruction (5).

Unlike most other pathogens, *Coxiella burnetii*, the causative agent of Q fever, subverts macrophage antimicrobial functions to establish a replicative niche termed the *Coxiella*-containing vacuole (CCV) that matures by fusing with lysosomal, autophagic, and secretory vesicles (6–8). Q fever is a worldwide zoonosis with significant public health implications (9). *C. burnetii* infects mammals (mainly cattle, goats, and sheep) and is typically transmitted to humans by aerosols derived from infected animal birth products (10). *C. burnetii* has been isolated from a variety of hosts and geographic regions (1, 11, 12). The original isolate, *C. burnetii* Nine Mile I (NMI), causes acute Q fever, whereas several others have been linked to chronic Q fever, including G (Q212), which was isolated from the heart valve of a person with endocarditis.

In contrast to strains that are pathogenic to humans, Dugway isolates collected from rodents have attenuated virulence in animal models and are not known to cause Q fever in humans (13). Because clinical and environmental isolates of *C. burnetii* require biosafety level 3 (BSL-3) containment, most laboratories use the BSL-2 strain *C. burnetii* Nine Mile II (NMII) to study *C. burnetii*-host interactions. NMII was derived from NMI through serial laboratory passage in embryonated eggs and has lost several lipopolysaccharide biosynthesis genes, which has rendered it avirulent (14).

In this project, to better understand macrophage response to *C. burnetii*, we investigated the transcriptomes of human alveolar macrophages (hAMs) infected with *C. burnetii* strains. We validated the gene expression data by measuring cytokine production and performing single-cell RNA-sequencing (scRNA-seq) of THP-1 macrophages infected with NMII. Our data show that proinflammatory pathways, including IL-17 signaling, are activated in *C. burnetii*-infected macrophages, but the inflammatory response varies depending on the bacterial strain and the macrophage subpopulation.

## RESULTS

### Macrophage gene expression varied between *C. burnetii* infections

We infected hAMs with *C. burnetii* NMI, NMII, G (Q212), or Dugway and measured gene expression at 72 hours post-infection (hpi) using RNA sequencing (RNA-seq). This analysis identified hundreds of genes that were differentially expressed in infected macrophages compared to uninfected controls (log2fc ≥ 0.75, padj < 0.05; n = 3) (**Figure 1, Table S1**). Infections with NMI or G (Q212), which are pathogenic to humans, or with NMII, a non-pathogenic strain derived from NMI, resulted in differential expression of roughly the same number of genes: 250, 266, and 332, respectively. In contrast, Dugway, an avirulent strain isolated from rodents, caused the differential expression of 672 genes, with most of the differentially expressed genes (445) showing down-regulation in infected cells compared to uninfected controls. Thirty-four differentially expressed genes (DEGs) were common to all four infections, with majority being inflammation-related genes.

**Figure 1.**
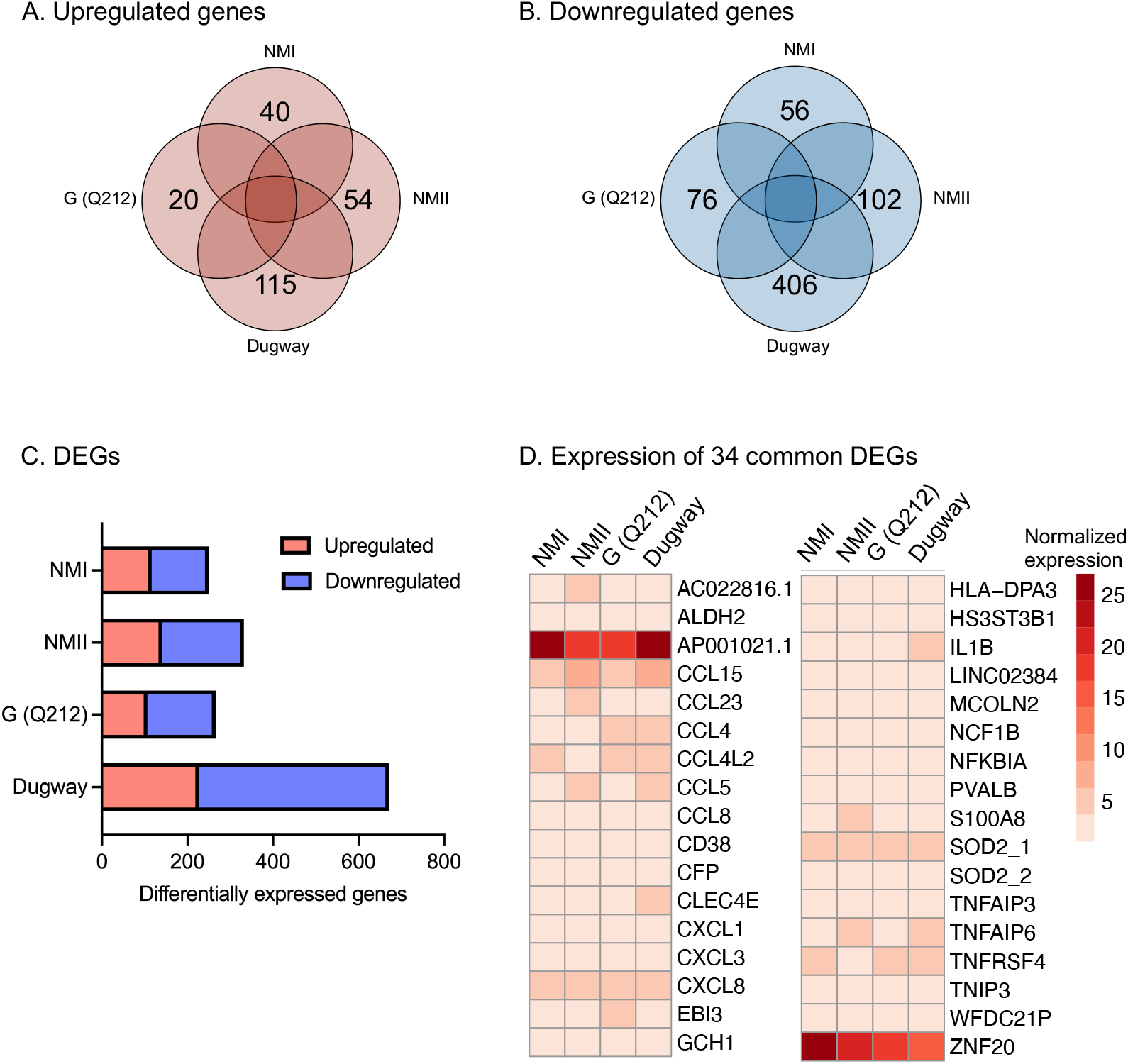
hAMs gene expression in response to *C. burnetii* infection. Venn diagrams showing (**A**) upregulated and (**B**) downregulated genes in primary human alveolar macrophages (hAMs) infected with NMI, NMII, G (Q212), or Dugway isolates of *C. burnetii* compared to uninfected cells (−0.75 ≤ log2fc ≥ 0.75, padj ≤ 0.05; n = 3). (**C**) Distribution of up- and down-regulated genes within differentially expressed genes (DEGs) in hAMs infected with each *C. burnetii* strain. (**D**) Heatmap of DEGs common to all four infections.

### *C. burnetii* activates macrophage proinflammatory pathways

To elucidate the functional implications of differential gene expression, we performed pathway enrichment analysis using the Ingenuity Pathway Analysis (IPA) software package (15). The IPA data showed that proinflammatory signaling, including IL-17 signaling, IL-6 signaling and TREM1 signaling, were activated (z-score ≥ 1.5, p < 0.05) in hAMs irrespective of the infecting *C. burnetii* isolate **(Figure 2, Table S2)**. IL-17, a member of the IL-17 family of proinflammatory cytokines, induces the expression of proinflammatory cytokines or chemokines in many cell types (16). To identify IL-17 signaling-associated proteins that may participate in *C. burnetii* infection, we used IPA to perform a pathway reconstruction analysis. We found that genes downstream of IL-17 signaling were consistently upregulated in *C. burnetii*-infected hAMs **(Figure 3, Table S1)**. The upregulated genes encode chemokines and cytokines that recruit immune cells or mount proinflammatory host response at the site of intracellular bacterial infection (17–23).

**Figure 2.**
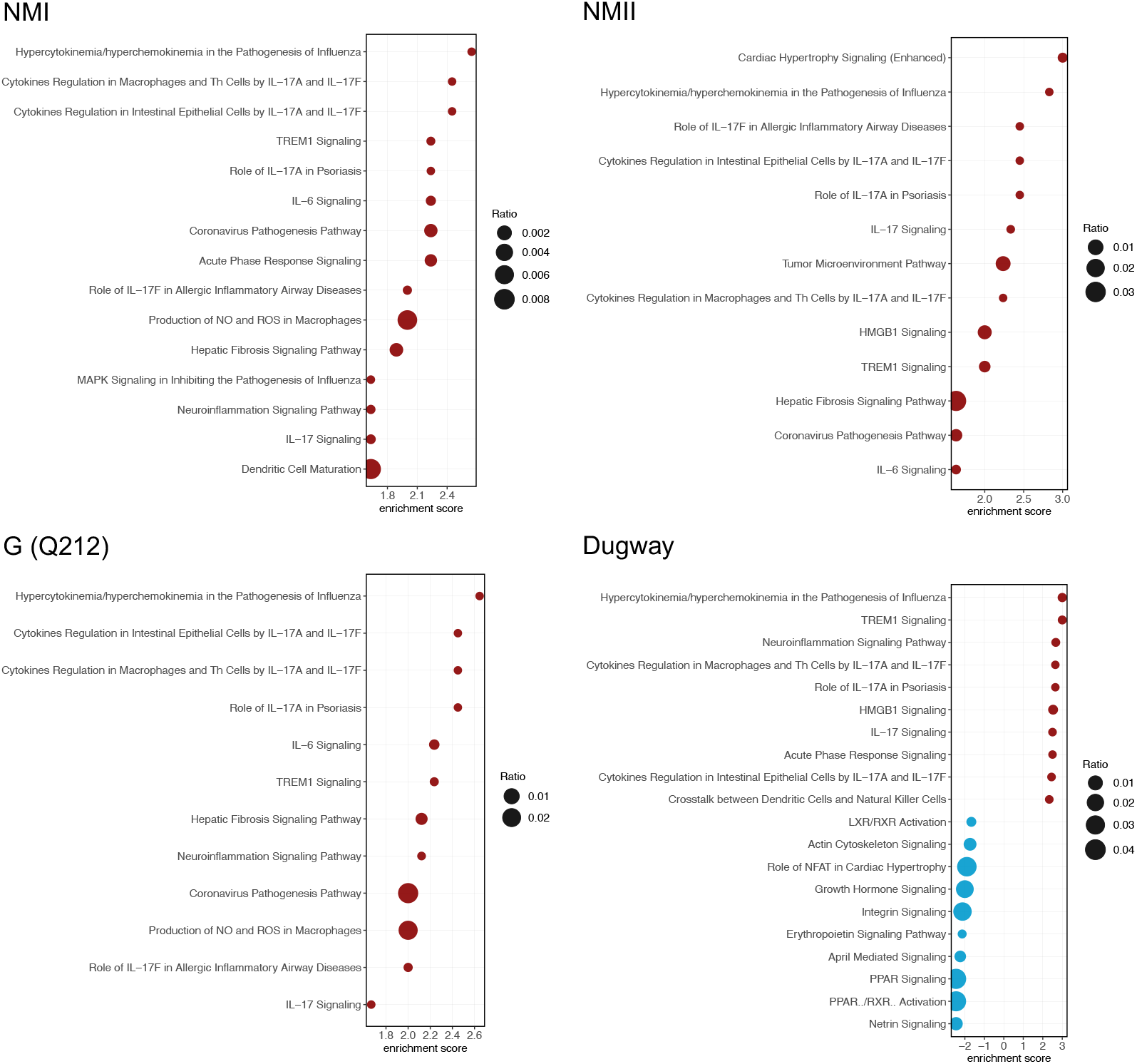
*C. burnetii* strains activate IL-17 signaling pathways. Bubble plots of top 20 enriched pathways in each infection. Red bubbles represent pathways that are activated (z-score ≥ 1.5, p<0.05) and blue bubbles correspond to inhibited pathways (z-score ≤ −1.5, p<0.05). Bubble size corresponds to the ratio of the number of differentially expressed genes in a pathway to the total number of genes in that pathway.

**Figure 3.**
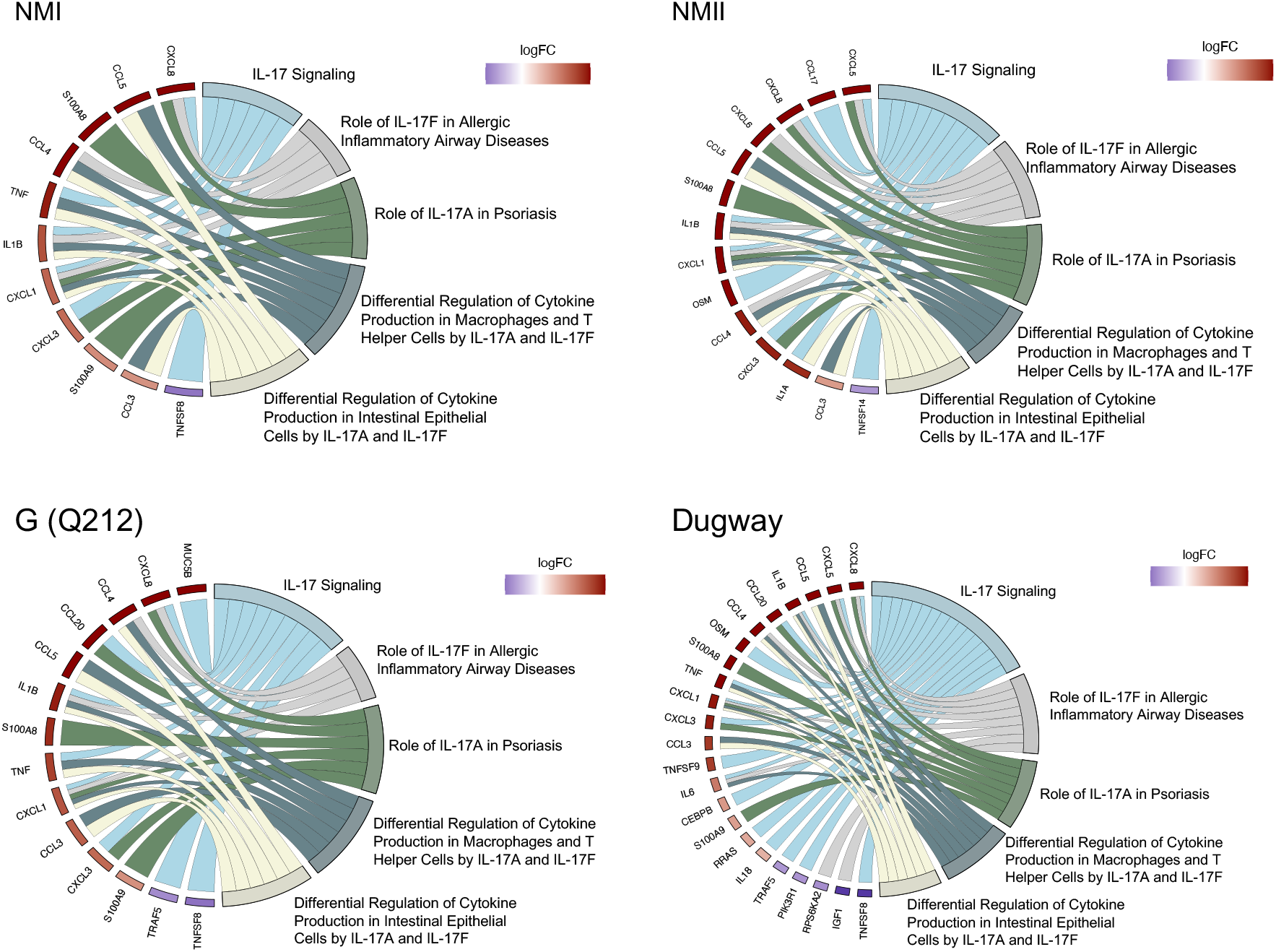
*C. burnetii* strains activate IL-17 signaling. Chord diagrams showing differential expression of genes (left) belonging to IL-17 signaling-related pathways (right) in each infection.

To validate the gene expression data, we infected THP-1 macrophages with NMII and measured several cytokines that are downstream of IL-17 signaling at 48 h, 72h, and 120 hpi (**Figure 4, Table S3**). This assay showed that significantly higher amounts of IL-1β, IL-8, GROα (CXCL1), CXCL10, CCL2, CCL3, CCL4 and CCL5 were secreted by infected macrophages at 120 hpi. But at 72 hpi only CXCL10, CCL3 and CCL4 and at 48 hpi only CSF2 and CXCL10 were produced at significantly higher amounts in infected cells (**Figure 4, Table S3**). These data suggest that immune response downstream of IL-17 signaling is not active in human macrophages during the initial stages of *C. burnetii* infection, as shown previously in mouse macrophages (24, 25), but by 120 hpi the proinflammatory IL-17 response is functional.

**Figure 4.**
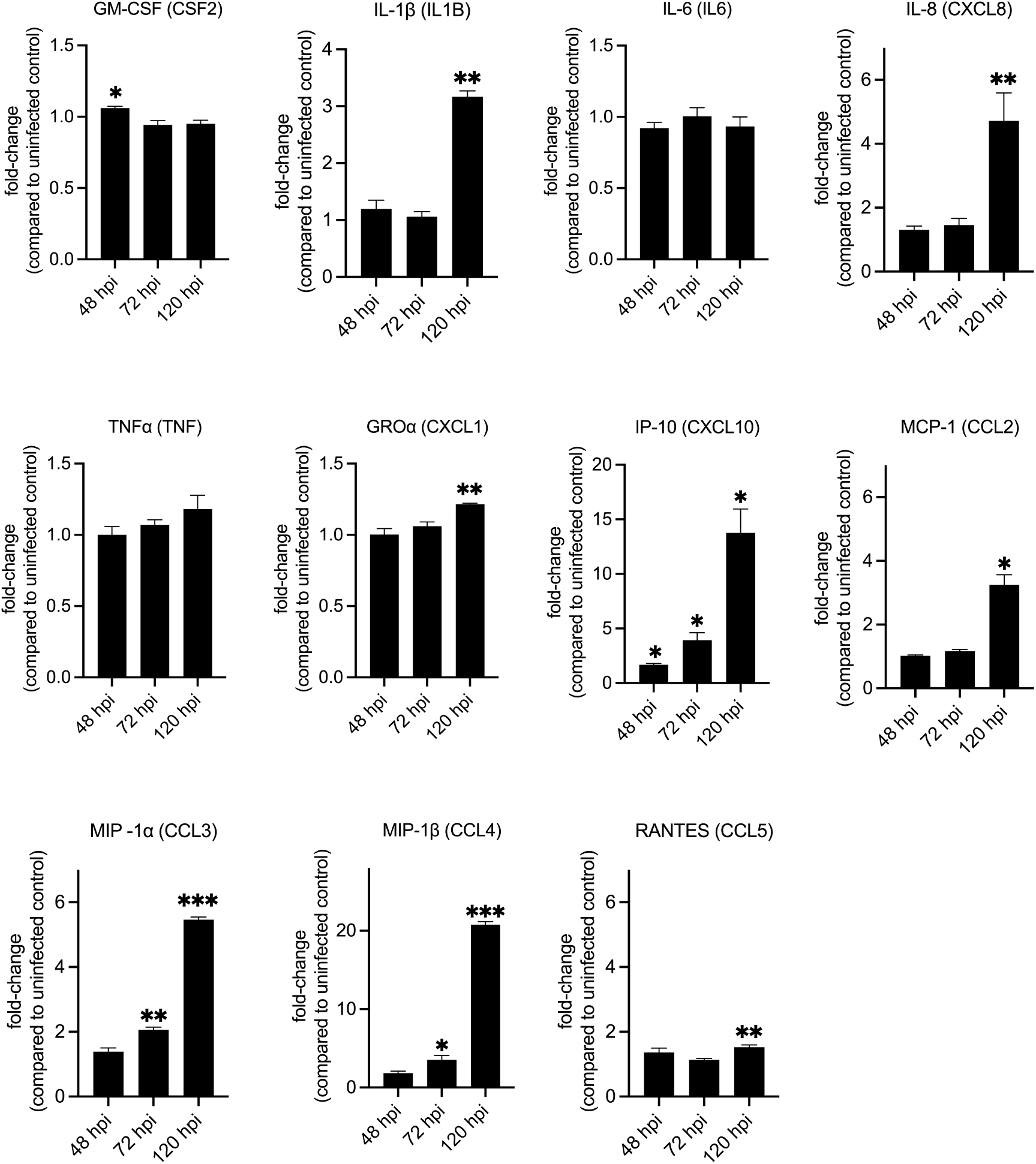
Secretion of cytokines downstream of IL-17 signaling in infected macrophages. Quantification of cytokines and chemokines released from NMII-infected THP-1 macrophages at 48, 72, and 120 h post-infection. Expression values are depicted as fold change between infected over uninfected cells at each timepoint. Statistical significance was calculated using two-tailed paired Student’s t-test, followed by Welch’s correction (*p < 0.05, **p < 0.01, ***p < 0.001, ****p < 0.0001; n = 3).

### Macrophage polarization varied between *C. burnetii* isolates

Using MacSpectrum (26), a tool that assigns macrophage polarization index (MPI) based on the expression of a set of genes that are associated with either M1 or M2 activation states, we estimated the polarization states of hAMs infected with NMI, NMII, G (Q212) or Dugway (**Figure 5)**. A high MPI indicates an M1-like (more inflammatory) phenotype, whereas a low MPI suggests an M2-like (less inflammatory) phenotype. The three uninfected samples had low MPI (−17.20, −10.03, −2.43), confirming that the inflammatory pathways in the control cells were not activated. In contrast, the three transcriptomes of Dugway-infected hAMs had high MPI (4.50, 13.74, 17.57), indicating that the rodent isolate induced a strong M1-like proinflammatory response in human macrophages. Transcriptional profiles of G (Q212)-infected hAMs also received positive MPI — albeit lower values than Dugway infection — (0.20, 1.73, 9.49). Unlike the MPIs for Dugway and G infections, which were relatively consistent across hAMs derived from three donors, infections with NMI and NMII produced a broad range of MPI (NMI: −7.39, - 1.94, 13.53; NMII: −18.29, 7.09, 8.83) (**Figure 5**). This wide distribution of MPI suggests that the immune response to NMI and NMII vary considerably between macrophage populations.

**Figure 5.**
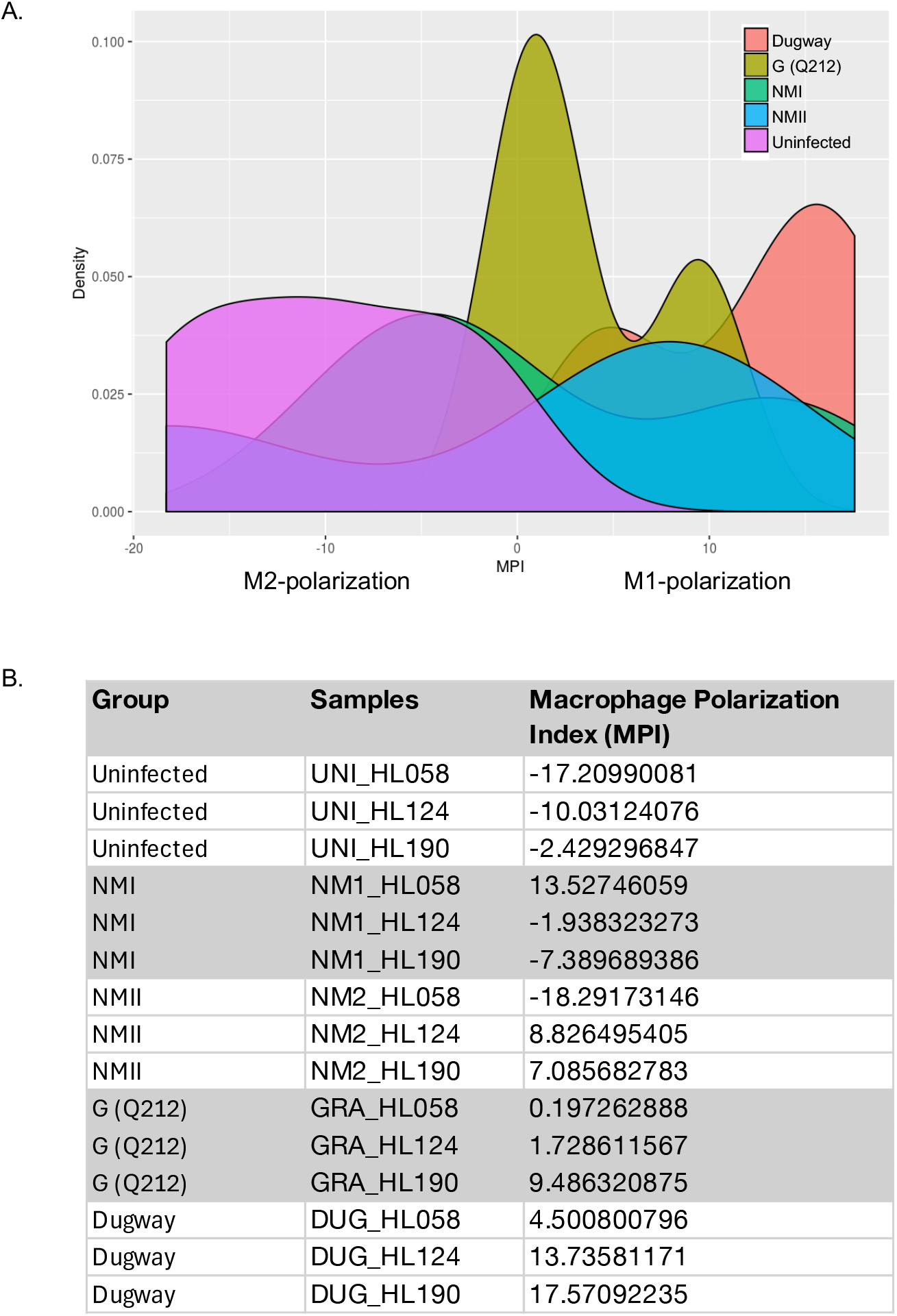
Macrophage polarization. (**A**) Density plots of Macrophage Polarization Index (MPI) for hAMs infected NMI, NMII, G (Q212), or Dugway isolates of *C. burnetii*. MPI were calculated using MacSpectrum and normalized gene expression values. (**B**) Table listing MPI for each sample. hAMs were derived from three donors (HL058, HL124, HL190).

### Single-cell sequencing show heterogenous macrophage response to NMII infection

To study macrophage inflammatory response to *C. burnetii* in more detail, we conducted a single-cell RNA-seq (scRNA-seq) analysis. We infected THP-1 macrophages with GFP-tagged NMII (27) and at 48 hpi sorted the cells into GFP-positive (NMII-infected) and GFP-negative (bystander) populations. Uninfected THP-1 macrophages that were processed similarly were used as controls (**Figure 6**). Around 1000 cells from each population were subjected to scRNA-seq, which revealed clusters (subpopulations) of cells with differing gene expression patterns within each population (**Figure 7**).

**Figure 6.**
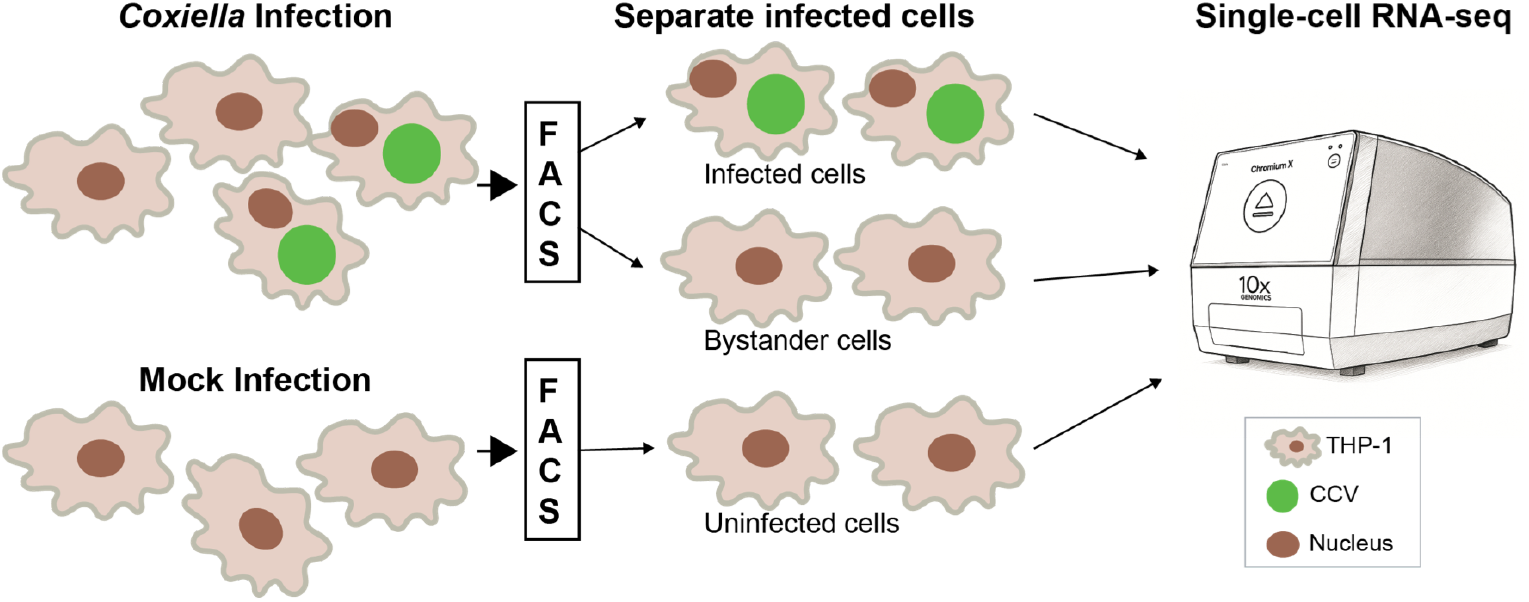
Single-cell RNA-sequencing workflow. THP-1 macrophages infected with GFP-tagged NMII were separated at 48 hours post-infection using fluorescence-activated cell sorting (FACS) into infected cells (GFP-positive) and bystander cells (GFP-negative). Mock-infected cells that were passed through the cell sorter were used as controls. Each population was processed using the 10x Genomics Chromium platform for single-cell RNA-sequencing. CCV, *Coxiella*-containing vacuole.

**Figure 7.**
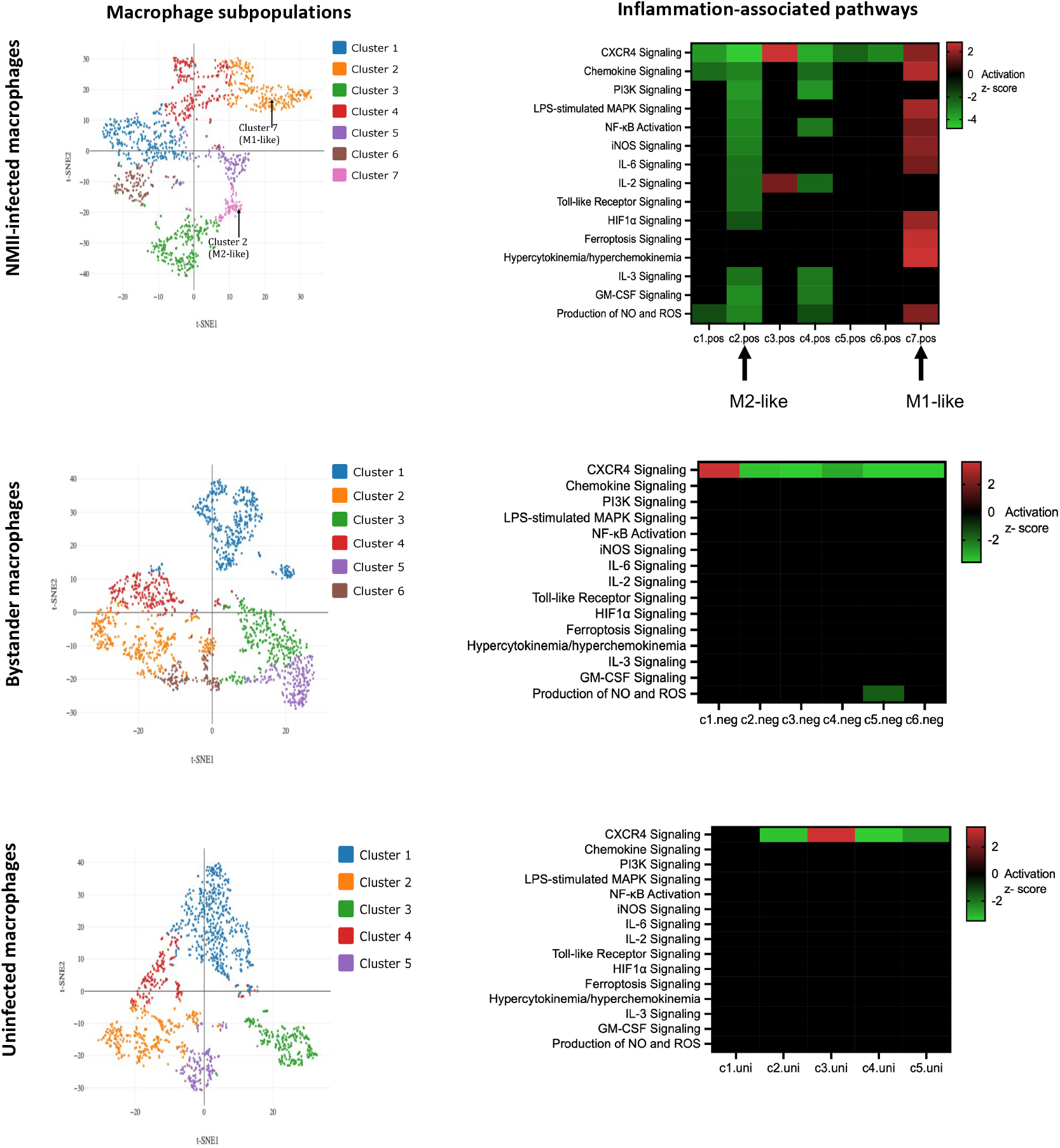
Single-cell analyses show transcriptional heterogeneity. (**Left panels**): Clustering of cells by t-distributed stochastic neighbor embedding (t-SNE) in NMII-infected (GFP-positive), bystander (GFP-negative) and uninfected THP-1 cells. (**Right panels**): Heatmaps depicting enrichment of 15 proinflammatory pathways (y-axes). Clusters within each population is shown on x-axes. Upregulated pathways are in red and downregulated pathways are in green (z-score ≥ 1.5 or ≤ −1.5, p < 0.05).

Pathway enrichment analysis of differentially expressed genes in NMII-infected macrophage subpopulations showed significant difference in the activation states of several inflammation-associated pathways **(Figure 7)**. For example, among 15 inflammation-associated signaling pathways (15, 28), 10 proinflammatory pathways, including pathways that are downstream to IL-17 signaling such as IL-6, MAPK, NFKB, iNOS, and NO/ROS signaling were significantly activated (positive z-score) in Cluster 7, whereas the same pathways were inhibited (negative z-score) in Cluster 2. Thus, cells in Cluster 7 appears to show a proinflammatory M1-like macrophage polarization, whereas cells in Cluster 2 have an anti-inflammatory M2-like polarization (29). Unlike in the NMII-infected macrophages, M1/M2-like polarization was not apparent in bystander or uninfected populations.

## DISCUSSION

In this study we show that *C. burnetii* activates proinflammatory pathways, particularly IL-17 signaling, in human macrophages. Interestingly, inflammatory response downstream of IL-17 signaling seemed to be muted during initial stages of *C. burnetii* infection and the magnitude and nature of the inflammatory response varied depending on the bacterial strain and the macrophage subpopulation.

*C. burnetii* Dugway, a rodent-derived avirulent strain, induced a robust transcriptional response in hAMs, including strong activation of proinflammatory pathways and consistent M1-like polarization. In contrast, the human-pathogenic isolates NMI and G (Q212), as well as the attenuated NMII strain, elicited a more subdued and variable response. The strong inflammatory response to Dugway, which likely contributes to its avirulence, suggests that the rodent isolate is less adapted to infecting human macrophages than NMI and G (Q212). Interestingly, Dugway infection also caused the downregulation of several signaling pathways. The processes that were downregulated only in Dugway infections include suppression of PPAR/RXR signaling that could impair lipid metabolism, and inhibition of integrin signaling that could disrupt cytoskeletal dynamics. At present it is unknown how, or if, these pathways contribute to the attenuated virulence of Dugway but identifying host processes that reduce C. *burnetii* virulence could contribute to the development of novel host-targeted therapies.

In our data, IL-17 signaling emerged as a central node in macrophage response to *C. burnetii*. This is not unexpected because IL-17 signaling is known to provide a protective immune response against other intracellular pathogens such as *L. pneumophila, M. tuberculosis*, and *Francisella tularensis* (3, 4, 30). IL-17 is a proinflammatory cytokine that activates several independent signaling mechanisms via TRAF (TNF receptor associated factor) and PI3K/AKT signaling (17, 31). These signaling cascades, in turn, lead to the secretion of chemokines and cytokines. Our transcriptomic and cytokine data for IL-17 signaling showed a temporal pattern, with significant gene expression occurring at 72 hpi but significant cytokine secretion occurring only at 120 hpi. The lag in cytokine secretion may reflect an initial suppression of IL-17 signaling pathway by *C. burnetii*, likely mediated via T4SS effectors, as observed in murine alveolar macrophages (24).

Macrophage polarization plays a critical role in shaping the immune response to intracellular pathogens. We observed that Dugway- and G (Q212)-infected hAMs consistently exhibited an M1-like phenotype, characterized by high macrophage polarization index (MPI). In contrast, NMI and NMII infections produced MPI scores that ranged from M2-like to M1-like in hAMs derived from different donors, suggesting that host genetic or epigenetic factors could influence polarization outcomes. This variability was also observed in the scRNA-seq data, which revealed distinct macrophage subpopulations with divergent polarization states in NMII-infected THP-1 macrophages. These findings underscore the importance of cellular heterogeneity in host-pathogen interactions and highlight the limitations of bulk transcriptomic analyses, which may obscure critical subpopulation dynamics.

Besides the induction of proinflammatory signaling, our gene expression analyses identified the activation of several pro-survival pathways, including PI3K/Akt signaling, autophagy, toll-like receptor signaling, TGF-β signaling, JAK/STAT signaling, STAT3 pathway, and MAPK signaling in *C. burnetii* infected hAMs. Our data also revealed putative roles for Wnt/Ca+ pathway and ferroptosis signaling that were previously unknown in the context of *C. burnetii* infection but are involved in infections with other pathogens such as *Ehrlichia chaffeensis* and *M. tuberculosis* (32, 33). Going forward, pursuing functional investigations of these pathways would likely advance our understanding of host-*C. burnetii* interactions.

## MATERIALS AND METHODS

### RNA sequencing and pathway analysis

Primary human alveolar macrophages (hAMs) were harvested by bronchoalveolar lavage (BAL) from postmortem human lung donors, as previously described (34). hAMs were infected with *C. burnetii* Nine Mile phase I RSA493 (NMI), Nine Mile phase II RSA439 (NMII), Dugway (5J108-111), or G (Q212) isolates at a multiplicity of infection (MOI) of 25. Infected and uninfected cells were cultured at 37°C under 5% CO_2_ in Dulbecco’s modified Eagle/F-12 (DMEM/F12) medium (Gibco) containing 10% FBS for 72 h. At this time-point, the growth medium was replaced with 1ml of TRI reagent (Life Technologies), and total RNA was extracted, and DNase treated (Invitrogen) as per manufacturer’s instructions. Samples were sequenced using the Illumina NovaSeq 6000 platform at Yale Center for Genome Analysis. Differential gene expression analysis was performed by mapping the sequencing reads to the reference human genome (GRCh38) using CLC Genomics Workbench v6.5 (Qiagen) and DESeq2 (35). Differentially regulated genes were calculated using log2 fold-change ≥ 0.75 and an adjusted p-value < 0.05 as cutoffs, as we described previously (36). Core analysis of the differentially expressed genes to find the enriched pathways was performed using Ingenuity Pathway Analysis (IPA, Qiagen) (15). For pathway analysis, gene clusters were compared with a standard mammal background database using z-score *≥* 1.5 or ≤ −1.5 (p < 0.05). Normalized counts from the RNA-seq data were used in MacSpectrum to determine macrophage polarization states (26).

### Cytokine secretion assay

*C. burnetii* NMII was cultured in acidified citrate cysteine medium-2 (ACCM-2) for 7 days at 37°C, 5% CO_2_, 2.5% O_2_ (37). Bacteria were quantified using PicoGreen (38), concentrated by centrifugation (3000xg, 10min, 4°c), and resuspended in PBS containing 0.25 M sucrose (PBSS) and stored at –80°C until further use. Before infection, THP-1 cells (American Type Culture Collection, TIB-202) were differentiated in RPMI-1640 medium supplemented with 1 mM sodium pyruvate, 0.05 mM beta-mercaptoethanol, 4500 mg/L glucose, and 10% heat-inactivated fetal bovine serum (FBS) at 37°C under 5% CO_2_ for 24 h using 30 nM phorbol 12-myristate 13-acetate (PMA), followed by 24 h of rest in PMA-free medium. THP-1 macrophages were infected with NMII at an MOI of 25. As positive control for the cytokine assay, THP-1 cells were treated with 200 ng/mL of *E. coli* O26:B6 lipopolysaccharide (Sigma-Aldrich) for 3h and 5 mM ATP (Adenosine 5′-triphosphate disodium salt hydrate; Sigma-Aldrich) for 30 min. Cytokine levels were assessed in cell culture supernatants using a Th1/Th2 Cytokine & Chemokine 20-Plex ProcartaPlex Panel 1 (Invitrogen) at 48, 72, and 120 hpi according to manufacturer’s guidelines. Briefly, supernatants were centrifuged 10,000xg for 10 min to remove particulate matter and stored at −80°C till further use. In a 96-well plate, magnetic beads were added in appropriate wells, washed, and 50µL of prepared antigen standards, controls, or samples were added. The plate was shaken for 30 min, 500 rpm at room temperature (RT), followed by overnight incubation at 4°C. The plate was shaken again for 30 min at RT (500 rpm), washed with 1X Wash Buffer, and incubated with the detection antibody (30 min, RT, 500 rpm). The plate was washed with 1X Wash Buffer, incubated with Streptavidin-Phycoerythrin (SAPE; 30 min, RT, 500rpm), followed by washing and addition of 120 µL of Reading buffer to analyze median fluorescence intensity (MFI) using a Luminex 200 instrument.

### Single-cell RNA sequencing

PMA-differentiated THP-1 cells were infected with GFP-tagged NMII (Tn1832) (27) at an MOI of 25 and at 48 hpi fluorescence-activated cell sorting (BD FACSAria Fusion) was carried out to separate infected (GFP positive) cells from bystander (GFP negative) cells. Uninfected THP-1 cells were similarly processed to serve as control. Single-cell RNA sequencing (scRNA-seq) libraries were generated at Yale Center for Genome Analysis from at least 1000 cells from each population by capturing individual cells inside gel beads in emulsion using single cell 3’ v3 chemistry (10X Genomics). Single-cell sequencing reads were processed for quality control and analyzed to compare cell clustering and differential gene expression within each population using Cell Ranger 3.1 pipeline (39). Single cell clusters were visualized using tSNE analysis using Loupe Cell Browser (10X Genomics) and log2 fold change was defined as the ratio of UMI (unique molecular identifier) counts in each cluster relative to all other clusters (log2 fold change ≥ 0.75 and p-value < 0.05). Differentially regulated genes in each cluster were analyzed using IPA to identify enriched pathways.

## DATA AVAILABILITY

Sequencing reads from this study have been deposited at NCBI Sequence Read Archive (SRA) under the BioProject accession PRJNA679931.

## ACKNOWLEDGEMENTS

This work was supported by funds from University of Texas at San Antonio and by National Institutes of Health grants AI123464 and AI133023.

## SUPPLEMENTARY TABLES

**Table S1**. Differentially expressed genes in hAMs infected with *C. burnetii* strains.

**Table S2**. Enriched pathways in hAMs infected with *C. burnetii* strains.

**Table S3**. Cytokine secretion assay.

## REFERENCES

1. Graham JG, MacDonald LJ, Hussain SK, Sharma UM, Kurten RC, Voth DE. 2013. Virulent Coxiella burnetii pathotypes productively infect primary human alveolar macrophages. Cell Microbiol 15:1012–1025.

2. Marriott HM, Dockrell DH. 2007. The role of the macrophage in lung disease mediated by bacteria. Exp Lung Res 33:493–505.

3. Gopal R, Monin L, Slight S, Uche U, Blanchard E, Fallert Junecko BA, Ramos-Payan R, Stallings CL, Reinhart TA, Kolls JK, Kaushal D, Nagarajan U, Rangel-Moreno J, Khader SA. 2014. Unexpected role for IL-17 in protective immunity against hypervirulent Mycobacterium tuberculosis HN878 infection. PLoS Pathog 10:e1004099.

4. Kimizuka Y, Kimura S, Saga T, Ishii M, Hasegawa N, Betsuyaku T, Iwakura Y, Tateda K, Yamaguchi K. 2012. Roles of interleukin-17 in an experimental Legionella pneumophila pneumonia model. Infect Immun 80:1121–1127.

5. Sun L, Wang L, Moore BB, Zhang S, Xiao P, Decker AM, Wang H-L. 2023. IL-17: Balancing Protective Immunity and Pathogenesis. J Immunol Res 2023:3360310.

6. Larson CL, Martinez E, Beare PA, Jeffrey B, Heinzen RA, Bonazzi M. 2016. Right on Q: genetics begin to unravel Coxiella burnetii host cell interactions. Future Microbiol 11:919– 939.

7. van Schaik EJ, Chen C, Mertens K, Weber MM, Samuel JE. 2013. Molecular pathogenesis of the obligate intracellular bacterium Coxiella burnetii. Nat Rev Micro 11:561–573.

8. Voth DE, Heinzen RA. 2007. Lounging in a lysosome: the intracellular lifestyle of Coxiella burnetii. Cell Microbiol 9:829–840.

9. Tan T, Heller J, Firestone S, Stevenson M, Wiethoelter A. 2024. A systematic review of global Q fever outbreaks. One Health 18:100667.

10. Van den Brom R, van Engelen E, Roest HIJ, van der Hoek W, Vellema P. 2015. Coxiella burnetii infections in sheep or goats: an opinionated review. Vet Microbiol 181:119–129.

11. Beare PA, Unsworth N, Andoh M, Voth DE, Omsland A, Gilk SD, Williams KP, Sobral BW, Kupko JJ, Porcella SF, Samuel JE, Heinzen RA. 2009. Comparative genomics reveal extensive transposon-mediated genomic plasticity and diversity among potential effector proteins within the genus Coxiella. Infect Immun 77:642–656.

12. Samuel JE, Frazier ME, Mallavia LP. 1985. Correlation of plasmid type and disease caused by Coxiella burnetii. Infect Immun 49:775–779.

13. Tesfamariam M, Binette P, Cockrell D, Beare PA, Heinzen RA, Shaia C, Long CM. 2022. Characterization of Coxiella burnetii Dugway Strain Host-Pathogen Interactions In Vivo. Microorganisms 10.

14. Millar JA, Beare PA, Moses AS, Martens CA, Heinzen RA, Raghavan R. 2017. Whole-Genome Sequence of Coxiella burnetii Nine Mile RSA439 (Phase II, Clone 4), a Laboratory Workhorse Strain. Genome Announc 5.

15. Krämer A, Green J, Pollard J, Tugendreich S. 2014. Causal analysis approaches in Ingenuity Pathway Analysis. Bioinformatics 30:523–530.

16. McGeachy MJ, Cua DJ, Gaffen SL. 2019. The IL-17 Family of Cytokines in Health and Disease. Immunity 50:892–906.

17. Amatya N, Garg AV, Gaffen SL. 2017. IL-17 Signaling: The Yin and the Yang. Trends Immunol 38:310–322.

18. Raimondo A, Lembo S, Di Caprio R, Donnarumma G, Monfrecola G, Balato N, Ayala F, Balato A. 2017. Psoriatic cutaneous inflammation promotes human monocyte differentiation into active osteoclasts, facilitating bone damage. Eur J Immunol 47:1062– 1074.

19. Cheung PFY, Wong CK, Lam CWK. 2008. Molecular mechanisms of cytokine and chemokine release from eosinophils activated by IL-17A, IL-17F, and IL-23: implication for Th17 lymphocytes-mediated allergic inflammation. J Immunol 180:5625–5635.

20. Maertzdorf J, Osterhaus ADME, Verjans GMGM. 2002. IL-17 expression in human herpetic stromal keratitis: modulatory effects on chemokine production by corneal fibroblasts. J Immunol 169:5897–5903.

21. Qian Y, Kang Z, Liu C, Li X. 2010. IL-17 signaling in host defense and inflammatory diseases. Cell Mol Immunol 7:328–333.

22. Jones CE, Chan K. 2002. Interleukin-17 stimulates the expression of interleukin-8, growth-related oncogene-alpha, and granulocyte-colony-stimulating factor by human airway epithelial cells. Am J Respir Cell Mol Biol 26:748–753.

23. Chung Y, Yamazaki T, Kim B-S, Zhang Y, Reynolds JM, Martinez GJ, Chang SH, Lim H, Birkenbach M, Dong C. 2013. Epstein Barr virus-induced 3 (EBI3) together with IL-12 negatively regulates T helper 17-mediated immunity to Listeria monocytogenes infection. PLoS Pathog 9:e1003628.

24. Clemente TM, Mulye M, Justis AV, Nallandhighal S, Tran TM, Gilk SD. 2018. Coxiella burnetii Blocks Intracellular Interleukin-17 Signaling in Macrophages. Infect Immun 86.

25. Clemente TM, Augusto L, Angara RK, Gilk SD. 2023. Coxiella burnetii actively blocks IL-17-induced oxidative stress in macrophages. BioRxiv.

26. Li C, Menoret A, Farragher C, Ouyang Z, Bonin C, Holvoet P, Vella AT, Zhou B. 2019. Single cell transcriptomics based-MacSpectrum reveals novel macrophage activation signatures in diseases. JCI Insight 5.

27. Martinez E, Cantet F, Fava L, Norville I, Bonazzi M. 2014. Identification of OmpA, a Coxiella burnetii protein involved in host cell invasion, by multi-phenotypic high-content screening. PLoS Pathog 10:e1004013.

28. Chen L, Deng H, Cui H, Fang J, Zuo Z, Deng J, Li Y, Wang X, Zhao L. 2018. Inflammatory responses and inflammation-associated diseases in organs. Oncotarget 9:7204–7218.

29. Viola A, Munari F, Sánchez-Rodríguez R, Scolaro T, Castegna A. 2019. The metabolic signature of macrophage responses. Front Immunol 10:1462.

30. Lin Y, Ritchea S, Logar A, Slight S, Messmer M, Rangel-Moreno J, Guglani L, Alcorn JF, Strawbridge H, Park SM, Onishi R, Nyugen N, Walter MJ, Pociask D, Randall TD, Gaffen SL, Iwakura Y, Kolls JK, Khader SA. 2009. Interleukin-17 is required for T helper 1 cell immunity and host resistance to the intracellular pathogen Francisella tularensis. Immunity 31:799–810.

31. Gu F-M, Li Q-L, Gao Q, Jiang J-H, Zhu K, Huang X-Y, Pan J-F, Yan J, Hu J-H, Wang Z, Dai Z, Fan J, Zhou J. 2011. IL-17 induces AKT-dependent IL-6/JAK2/STAT3 activation and tumor progression in hepatocellular carcinoma. Mol Cancer 10:150.

32. Luo T, Dunphy PS, Lina TT, McBride JW. 2015. Ehrlichia chaffeensis Exploits Canonical and Noncanonical Host Wnt Signaling Pathways To Stimulate Phagocytosis and Promote Intracellular Survival. Infect Immun 84:686–700.

33. Amaral EP, Costa DL, Namasivayam S, Riteau N, Kamenyeva O, Mittereder L, Mayer-Barber KD, Andrade BB, Sher A. 2019. A major role for ferroptosis in Mycobacterium tuberculosis-induced cell death and tissue necrosis. J Exp Med 216:556–570.

34. Graham JG, Winchell CG, Kurten RC, Voth DE. 2016. Development of an Ex Vivo Tissue Platform To Study the Human Lung Response to Coxiella burnetii. Infect Immun 84:1438– 1445.

35. Love MI, Huber W, Anders S. 2014. Moderated estimation of fold change and dispersion for RNA-seq data with DESeq2. Genome Biol. 15(12):550.

36. Sachan M, Brann KR, Fullerton MS, Voth DE, Raghavan R. 2023. MicroRNAs Contribute to Host Response to Coxiella burnetii. Infect Immun. 91(1):e0019922.

37. Omsland A, Beare PA, Hill J, Cockrell DC, Howe D, Hansen B, Samuel JE, Heinzen RA. 2011. Isolation from animal tissue and genetic transformation of Coxiella burnetii are facilitated by an improved axenic growth medium. Appl Environ Microbiol 77:3720–3725.

38. Moses AS, Millar JA, Bonazzi M, Beare PA, Raghavan R. 2017. Horizontally Acquired Biosynthesis Genes Boost Coxiella burnetii’s Physiology. Front Cell Infect Microbiol 7:174.

39. Zheng GXY, Terry JM, Belgrader P, Ryvkin P, Bent ZW, Wilson R, Ziraldo SB, Wheeler TD, McDermott GP, Zhu J, Gregory MT, Shuga J, Montesclaros L, Underwood JG, Masquelier DA, Nishimura SY, Schnall-Levin M, Wyatt PW, Hindson CM, Bharadwaj R, Wong A, Ness KD, Beppu LW, Deeg HJ, McFarland C, Loeb KR, Valente WJ, Ericson NG, Stevens EA, Radich JP, Mikkelsen TS, Hindson BJ, Bielas JH. 2017. Massively parallel digital transcriptional profiling of single cells. Nat Commun 8:14049.

